# Glutamatergic modulation of auditory cortex connectivity with attentional brain networks in unpredictable perceptual environment

**DOI:** 10.1101/2019.12.20.884049

**Authors:** K. Kompus, V. Volehaugen, A. Craven, K. Specht

## Abstract

In a stable environment the brain can minimize processing required for sensory input by forming a predictive model of the surrounding world and suppressing neural response to predicted stimuli. Unpredicted stimuli lead to a prediction error signal propagation through the perceptual network, and resulting adjustment to the predictive model. The inter-regional plasticity which enables the model-building and model-adjustment is hypothesized to be mediated via glutamatergic receptors. While pharmacological challenge studies with glutamate receptor ligands have demonstrated impact on prediction-error indices, it is not clear how inter-individual differences in the glutamate system affect the prediction-error processing in non-medicated state. In the present study we examined 20 healthy young subjects with resting-state proton MRS spectroscopy to characterize glutamate+glutamine (rs-Glx) levels in their Heschl’s gyrus (HG), and related this to HG functional connectivity during a roving auditory oddball protocol. No rs-Glx effects were found within the frontotemporal prediction-error network. Larger rs-Glx signal was related to stronger connectivity between HG and bilateral inferior parietal lobule during unpredictable auditory stimulation. We also found effects of rs-Glx on the coherence of default mode network (DMN) and frontoparietal network (FPN) during unpredictable auditory stimulation. Our results demonstrate the importance of Glx in modulating long-range connections and wider networks in the brain during perceptual inference.

## Introduction

The underlying principle of processing sensory information is predictive modeling: forming a model of likely perceptual inputs, and generating an error signal in the case of a mismatch in order to update the model. EEG and MEG studies with auditory stimuli have shown that mismatch detection occurs within 200 ms from the deviant feature onset, and can be explained by a network of sources in the superior temporal plane and inferior frontal cortex [1–3]. The early prediction error signal associated with deviance detection is likely the first step in a reorienting sequence [4], followed by P3a ERP which has been linked to higher-level attentional processes involving parietal and frontal areas [5], and thereafter P3b which has been connected to context updating involving temporo-parietal areas [6]. fMRI studies showing parietal and frontal activations [7, 8] support the view that attention capture by the stimuli leads to engagement of wider brain networks. The engagement of frontoparietal attention networks has been found in fMRI studies of oddball processing [9]. Centrally, the role of the inferior parietal lobe in conjunction with intraparietal sulcus is important in initiating attention shifts based on the priority map which interacts with the sensory areas [10], thus the frontoparietal attention network engagement in oddball studies is likely.

The neurophysiological underpinnings of the auditory change detection have been related to glutamate, the major excitatory neurotransmitter. Glutamatergic neurotransmission has been suggested to implement the top-down predictions and carry information about the bottom-up prediction errors [3]. Consequently, individuals with high glutamate levels could have more efficient prediction error signaling, as suggested by a combined MRS-EEG study [11].

How the glutamate modulates the signaling between the prediction error generators distributed within the network consisting of superior temporal plane and inferior frontal regions, as well as with the wide-spread brain networks supporting attention is an important question. One line of evidence comes from pharmacological challenge studies. Ketamine, an NMDA-receptor antagonist, reduced inter-regional functional connectivity within the auditory network while not affecting local adaptation [3]. Further, this correlated with lower self-ratings of “control and cognition”, indicating wider effects in the brain, including hierarchically higher levels. A single-trial analysis of a roving MMN paradigm based on the Hierarchic Gaussian Filter model suggested that ketamine reduced higher-order prediction errors expressed in the time window of P3a, reflecting changes to volatility estimates of stimulus transition probabilities due to NMDA receptor blockage [12]. A possible interpretation is that dysfunctional NMDA receptors led to overestimation of volatility, and thereby altered learning rate about the statistical structure of the environment: the individual would be less surprised by the changes because they would expect these to happen in a volatile environment. Correspondingly it could be hypothesized that the glutamatergic signal between the auditory and hierarchically higher areas supports the correct estimation of the higher-level statistical structure, and thereby allow efficient detection of surprising events that deviate from that structure. The other line of evidence comes from MRS-based studies of unperturbed glutamate. The MRS-measured rs-Glx in auditory cortex (composite of glutamate and glutamine measured during resting-state acquisition) has been shown to affect the processing of duration-deviants and gap-deviants [11] while leaving other deviants unaffected; an effect possibly linked to higher complexity, involving frontal sources, related to representing duration-and gap-predictions [13]. Taken together, glutamatergic signaling is clearly important for the information exchange between the auditory areas and higher-level, frontoparietal networks in order to represent the confidence in the environment in which the standards and deviants are embedded.

Another important question is the behavior of widespread brain networks during unpredictable auditory stimulation, and how this is modulated by glutamatergic neurotransmission. Ketamine has been shown to reduce deactivation in the task-negative default mode network and activation in the task-positive frontoparietal regions during a working memory task [14]. In agreement with this it has been shown that rs-Glu and GABA measured in posterior cingulate cortex predict task-induced deactivations in the entire (task-negative) default mode network during a working memory task [15], while rs-Glx in dorsal ACC predicted task-induced BOLD response in inferior parietal (task-positive) areas during high cognitive control demands [16]. Duncan and colleagues [17] conclude that the regional glutamate concentration effects are shown exclusively in inter-regional connections (outside the measured area), and hypothesize that the effects may reflect the long-range projections from glutamatergic excitatory neurons.

In the present study we examined how rs-Glx affects signaling in the frontotemporal auditory prediction error network, and relationship of the auditory cortex with wider networks (frontoparietal attention network and default mode network). We used a paradigm previously used to examine brain responses to auditory unpredictability in fMRI [8]. Subjects were exposed to blocks of tones in predictable order (ABAB) and unpredictable order (alternating A- and B- tone sequences of varying length, so-called ‘roving standard’ protocol). Two different deviating-feature types were used in separate conditions (frequency and duration). Glutamate was quantified from a resting-state measurement of Glx with ^1^H-MRS in Heschl’s gyrus (HG) before fMRI was used to measure the BOLD response to the auditory stimulation.

We hypothesized that:

1. during the unpredictable blocks individuals with higher rs-Glx show increased signaling in the frontotemporal auditory hierarchical prediction network consisting of bilateral HG, planum temporale (PT), posterior STG (pSTG) and inferior frontal gyrus (IFG) (Figure 1, right);
2. during the unpredictable blocks there should be increased connectivity between auditory cortex and areas involved in the frontoparietal attention network in individuals with higher rs-Glx;
3. rs-Glx levels relate to strength of the anticorrelation between task-negative (DMN) and task-positive (frontoparietal) networks during the unpredictable blocks, and increased within-network connectivity in the frontoparietal network.

**Figure 1.**
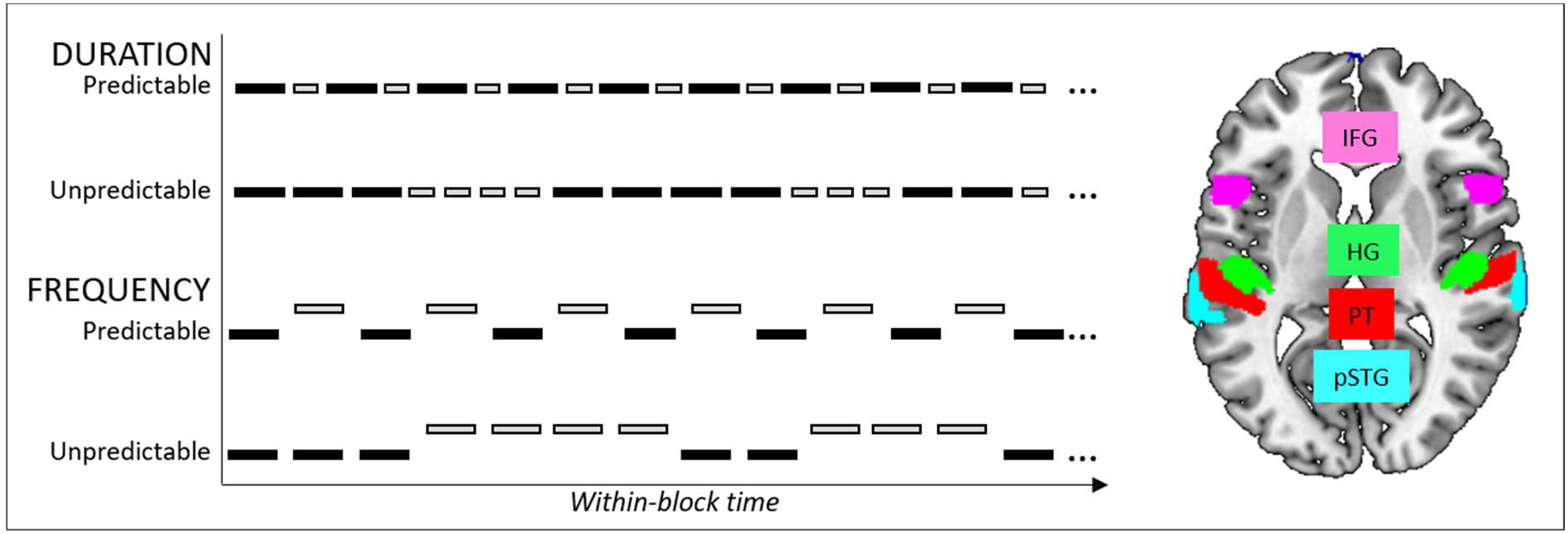
Left: Schematic presentation of the auditory stimulation protocol. The predictable and unpredictable sequences of A- and B-tones were presented within two conditions: duration and frequency. During the predictable blocks, the A- and B-tones alternated regularly, during the unpredictable blocks the trains of A- and B-tones alternated unpredictably. Right: The ROIs representing the frontotemporal prediction error network, including bilateral inferior frontal gyrus, pars opercularis (IFG), Heschl’s gyrus (HG), planum temporale (PT), posterior superior temporal gyrus (pSTG).

## Methods

### Subjects

The subjects were 20 healthy young right-handed adults (9 male, 11 female, age 20-38 years, mean 25.5, SD 4.8), psychiatrically and neurologically healthy by self-report. The subjects were asked to avoid consuming nicotine and caffeine within 5 hours before the experiment. The subjects’ hearing acuity was measured in both ears for frequencies 250, 500, 1000, 2000 and 3000 Hz. The first principal component of the hearing acuity was used as a covariate for all analyses.

### Auditory paradigm

The stimulation paradigm is presented in Figure 1 (left). The stimuli were composite tones, with the standard tone duration 100 ms (with 5 ms rise and fall time), composed of 8 equally loud sinusoidal tones (fundamental frequency 330 Hz, and seven harmonic partials). There were two conditions, order counterbalanced between the subjects: Duration (A tone: standard as described, B tone: length 60 ms) and Frequency (A tone: standard as described, B tone: 1.25 semitones higher). Within each condition, four predictable and four unpredictable blocks (length 25 s) were presented. During predictable blocks, A and B tone were alternating regularly (ABABAB…), during unpredictable blocks, A and B tone were alternating unpredictably (e.g., AABBBAAAABBA…); ISI was 320 ms. The subjects were instructed to ignore the tones while viewing a nature video.

### MR acquisition

The MR data were gathered on a 3T Siemens Prisma MR scanner. First, high-resolution anatomical images were acquired with T1-MPRAGE sequence, sequential increasing slice order, 192 slices, TR= 1800 ms, TE=2.28 ms, matrix = 256×256×192. The placement of the MRS voxel was determined on the anatomical image. The voxel placement covered the Heschl’s gyrus. The MRS acquisition was performed before the fMRI task in each subject. The order of the MRS acquisition was counterbalanced between the left and the right hemisphere. Short echo time water-suppressed 1H spectra were gathered with PRESS sequence (voxel dimensions 20*20*20 mm3, TE/TR = 30/2000 ms, 80 averages). Unsuppressed water reference spectra (4 averages) were acquired in the same voxel after the water-suppressed acquisition. The fMRI data was acquired as 320 images, matrix 64×64×35, TR 2000 ms, TE 30 ms, 35 slices. During the fMRI data acquisition, the auditory stimuli were presented.

### MRS data analysis

The MRS data were analyzed using LCModel (version 6.3-1L) (Provencher, 1993) with a standard basis set (Ala, Asp, Cr, PCr, GABA, Glc, Gln, Glu, GPC, PCh, GSH, Ins, Lac, NAA, NAAG, Scyllo, Tau), scaling the metabolite estimates to the internal water reference. For water-scaling, voxel-specific estimates calculated based on the tissue composition of the MRS voxel. Briefly, the estimates were corrected to account for the differences in water concentration and water relaxation times and partial volume effects depending on the tissue class inside the voxel (WM, GM, CSF). The fractional proportion of the GM, WM and CSF within the spectroscopy voxels was estimated using SPM12 routines to segment the anatomical image into grey matter, white matter and CSF images. The proportion of GM, WM and CSF was measured within the MRS voxel mask. The correction was implemented for each metabolite (term *metab*) according to the equation

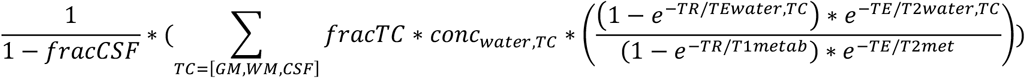

where frac indicates fraction of the tissue class in voxel, conc_water,TC_ is the concentration of water in the tissue class (conc GM/WM/CSF = 4330/36080/54840 mM respectively, Ernst 1993). The parameters for the metabolites of interest were: T1/T2glu: 1270/180 (Mlynarik et al., 2001, Ganji et al., 2012), T1/T2cre: 1460/152 (Mlynarik et al., 2001). Glx was expressed as creatine ratio (Glx/Cre) for consistency with previous reports on the relationship between Glx and neurocognitive variables [17]. To verify that significant effects are not due to creatine, we examined the significant connections in a regression analysis where we entered the water-scaled concentration of Glx and Cre separately, full details are reported in Supplementary Materials.

### fMRI data preprocessing

The preprocessing was conducted with SPM12 functions incorporated in CONN toolbox processing pipeline. The functional images were realigned, normalized to MNI space, and smoothed with a Gaussian kernel with 6 mm full width at half maximum. FSL Harvard-Oxford atlas was used to define the following ROIs: bilateral Heschl’s gyrus (L_HG and R_HG), bilateral pSTG (L_pSTG and R_pSTG), bilateral planum temporale (L_PT, R_PT) and bilateral inferior frontal gyrus, pars opercularis (L_IFG, R_IFG). The ROI data was extracted from non-smoothed volumes as the mean timeseries of the voxels within the ROI mask. Further preprocessing for the connectivity analysis consisted of removing the subject-specific movement parameters estimated during realignment step, the white matter and CSF signal, and the main effects of the task blocks (modeled as boxcar functions for the duration of the stimulation block, convolved with the hemodynamic response function), finally the data were filtered with a bandpass filter from 0.008 to 0.09 Hz.

Within each subject, the connectivity was calculated as bivariate correlation between the timeseries of all ROIs. For whole-brain seed-to-voxel analysis, the connectivity between the timeseries L_HG and R_HG and the rest of the voxels in the entire brain within each subject was calculated.

For the second-level analyses, we examined the change in the connectivity in both the seed-to-voxel and ROI-to-ROI analyses between the predictable and unpredictable blocks (unpred>pred contrast), as well as modulation of that change by rs-Glx. For this, the rs-Glx measures in the left and right hemisphere were entered into the second-level model as covariates.

The ROI-to-ROI analysis examined the effect of left and right rs-Glx on any of the connections between all the ROIs in the frontotemporal network during unpred>pred contrast (FDR-corrected at p<.05 for all connections in the analysis). Additionally we performed a planned test (one-sided uncorrected p<.05) for increased connectivity in the connections between L_HG and L_PT as well as L_HG and L_STG, due to earlier studies showing specific modulation of connection between HG and posterior STG by ketamine [3].

To explore connectivity changes due to rs-Glx outside of the frontotemporal network we performed a seed-to-voxel analysis, where we examined the effect of ipsilateral rs-Glx on the connectivity between each of the two HG ROIs and the rest of the brain during unpred>pred contrast (height threshold uncorrected p<.005 and cluster extent threshold FWE-corrected p<.01).

Finally, we tested whether the DMN and FPN change their properties depending on the predictability of the blocks, and whether this is modified by rs-Glx levels. The network anatomical definition was based on the CONN toolbox networks atlas. The DMN nodes consisted of medial PFC (centre x,y,z = 1,55,−3), bilateral parietal cortex (−39, −77, 33; 47, −67, 29) and posterior cingulate cortex (1, −61, 38). The FPN nodes consisted of bilateral prefrontal cortex (−43, 33, 28; 41, 38, 30) and posterior parietal cortex (−46, −58, 49; 52, −52, 45). We tested the within-network connectivity and between-network connectivity, defined as mean correlation across all nodes within each network, and mean correlation between all the nodes of two networks, respectively. The within- and between-network connectivity was calculated using the *withinbetweenROI* tool in the CONN toolbox. The statistical model, consisting of within-subject factors predictability and condition with between-subject covariates left rs-Glx and right rs-Glx as above was estimated using SPSS.

## Results

### ROI-to-ROI analysis: the effect of rs-Glx on the frontotemporal network

First, we examined the effect of predictability on the frontotemporal network (main effect of unpred>pred). While both during unpredictable and predictable blocks there was positive connectivity within the auditory network consisting of the temporal ROIs, this was lower in the unpredictable blocks (see Supplementary figure 1, left). Additionally, there was increased connectivity between L_IFG and L_HG; this was due to more positive-going connectivity in unpredictable blocks (Supplementary figure 1, right). There was no significant condition*predictability interaction, indicating that the observed connectivity changes due to unpredictability were common for Duration and Frequency conditions.

Next, we examined the effect of left and right rs-Glx on the frontotemporal network during unpred>pred. For the connectivity between all ROIs we did not find any effect of either left or right rs-Glx on the main effect of predictability, or the condition*predictability interaction.

The planned comparison concentrating on the connections between the L_HG and L_pSTG, L_PT did not show any rs-Glx effects on the main effect of predictability, or the condition*predictability interaction.

### Seed-to-voxel analysis: the effect of rs-Glx on whole-brain connectivity

For the whole-brain connectivity analysis of the unpred>pred effect with the seed in the L_HG we found an effect of left rs-Glx in bilateral inferior parietal lobule (MNI x, y, z = 50, −48, 32 and −40, −58, 42), see Figure 3. To verify that the effect was not due to creatine-scaling, we performed a hierarchical regression analysis with the water-scaled values of rs-Glx and rs-Cre, including the hearing acuity in the model. This control analysis (see Supplementary Materials) showed that water-scaled Glx was significantly related to the unpred>pred effect while controlling for the other variables (standardized beta = ․83, t=3.54, p=.003).

**Figure 2.**
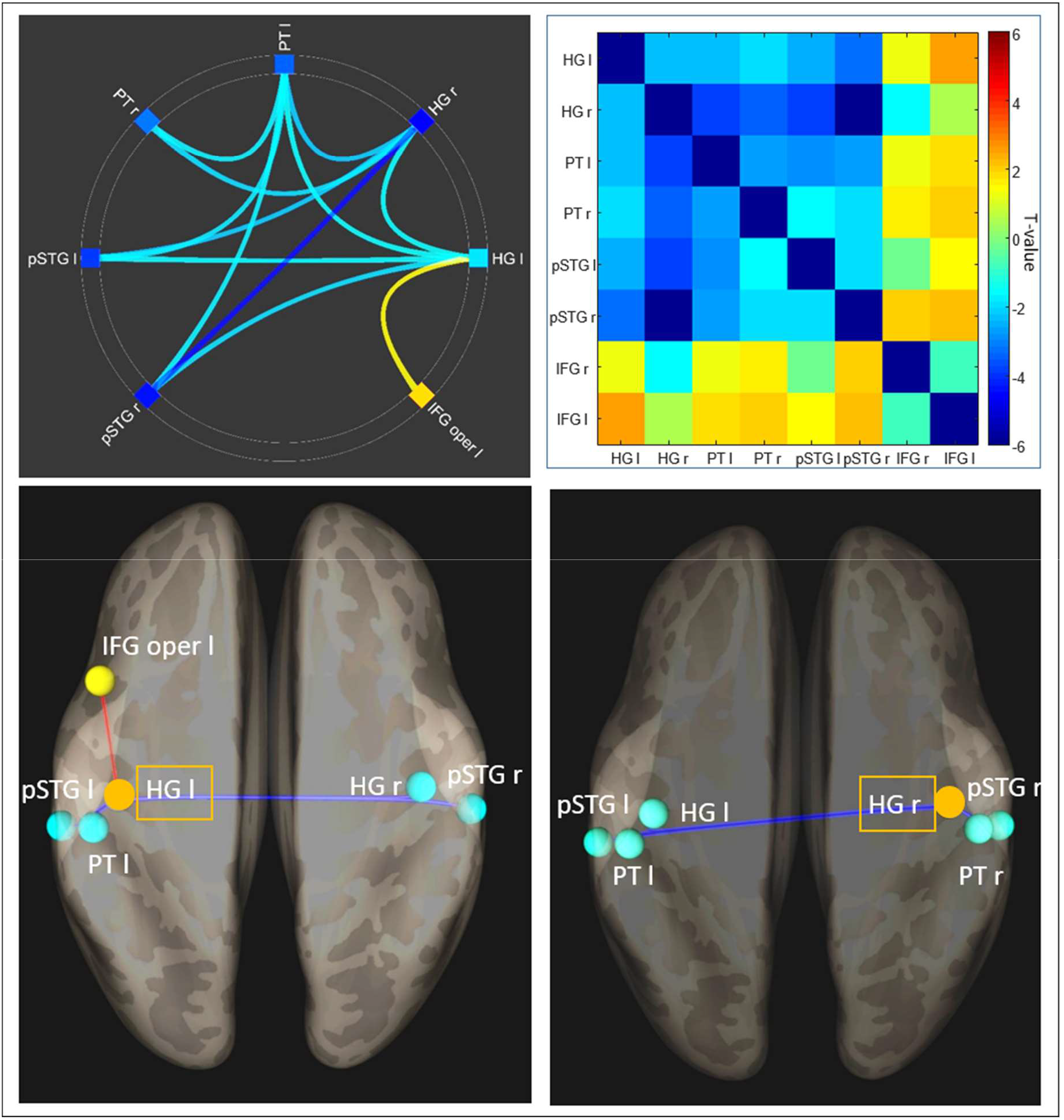
Top left: Unpred>pred main effect was significant in several connections (p<.05, two-sided, FDR-corrected across all connections in the analysis) in the frontotemporal network; we observed reduced connectivity during unpred blocks in the temporal areas, and increased connectivity during unpred blocks between left HG and left IFG. Top right: The full connectivity profile, including significant and non-significant results for the unpred>pred contrast, shows that overall the frontal-to-temporal connections were positive (increased connectivity during unpred blocks), while temporal-to-temporal connections were negative (reduced connectivity during unpred blocks). Bottom: The right and left HG connectivity profile was overall similar, with emphasis on the reduced connectivity during unpred blocks among the temporal areas. Left HG specifically showed increased connectivity with ipsilateral IFG during the unpred blocks.

**Figure 3.**
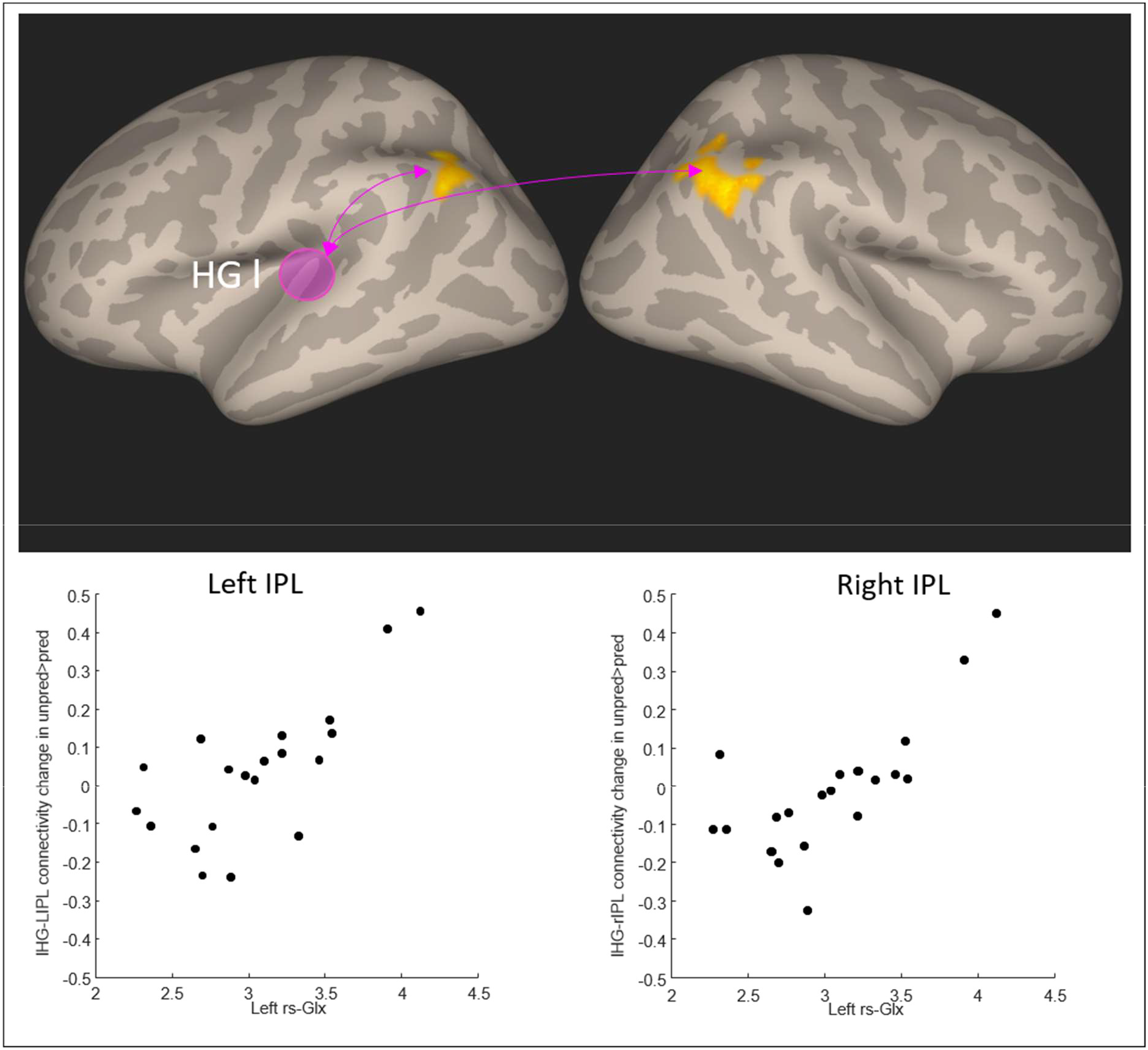
The connectivity between left HG (gyrus location in the figure indicated with the purple circle) and bilateral IPL during unpred blocks was modulated by the rs-Glx in the left HG. The scatterplots visualize the mean unpred>pred contrast value in the significant cluster in each subject against their left HG rs-Glx values.

**Figure 4.**
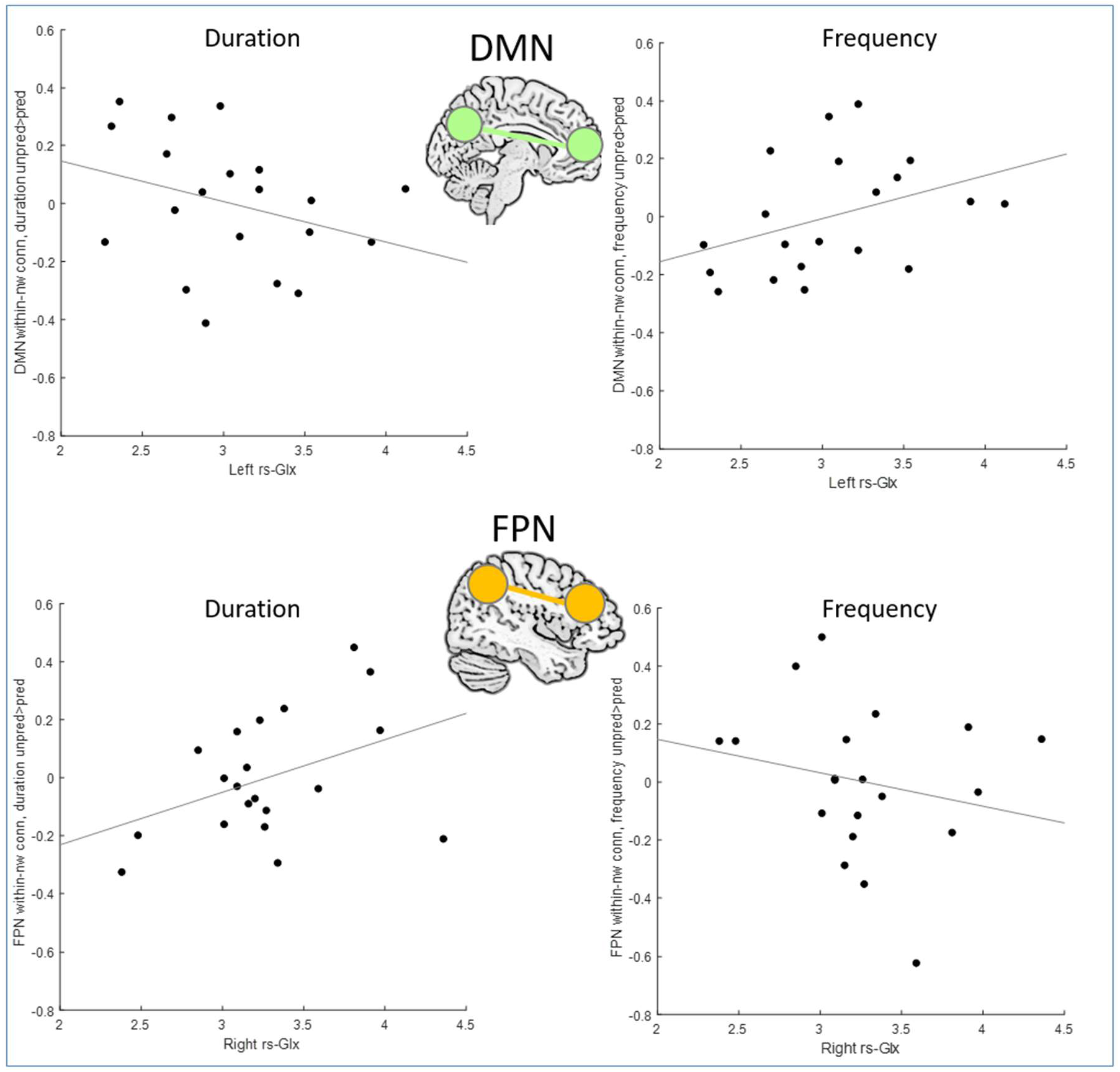
The within-network connectivity in DMN (top) and FPN (bottom) during unpred>pred contrast, separately for frequency (right) and duration (left), visualized in scatterplots against the rs-Glx in the HG, with least-squares regression line to illustrate the trend.

Using the SPM Anatomy toolbox [18] to localize the significant clusters showed that in the right hemisphere, the activation was in areas PGa (29.6 % of the anatomical area covered), PGp (14.4 % area covered), intraparietal sulcus area hIP1 (18 % area covered), and PFm (6 % area covered). In the left hemisphere the activation was in areas PGa (32.4 % area covered), PGp (7 % area covered), PFm (8.7 % area covered), and intraparietal sulcus area hIP1 (10.8 % area covered). We did not find any effects for the analysis in the right hemisphere (seed R_HG).

### The effect of rs-Glx on DMN and FPN within-network connectivity

We hypothesized that higher rs-Glx level should lead to larger unpred-pred difference in the connectivity within the examined networks (FPN and DMN), reflecting the brain’s increased ability to react to unpredictable stimulation by recruiting the FPN while disengaging the DMN.

In DMN, in all four conditions there was significant positive connectivity within the network (duration pred, duration unpred, frequency pred, frequency unpred: Cohen’s d respectively 2.56, 2.58, 2.17, 2.19). There was a small but significant effect of left rs-Glx on the interaction between feature type and deviation (p=.019, F(1,16)=6.75, η^2^=0.05). To explore this effect, we calculated the unpred>pred difference for both duration and frequency condition, and tested for the relationship between the unpred>pred difference and left rs-Glx. The post-hoc regression analysis, including hearing accuracy and right rs-Glx as covariates, showed that the effect of left rs-Glx on unpred>pred difference was different in the two conditions, being a weaker negative relationship in duration (standardized beta= −0.34, t= −1.47) and a stronger positive relationship in frequency (standardized beta= 0.41, t= 1.81).

In FPN, in all four conditions there was significant positive connectivity within the network (duration pred, duration unpred, frequency pred, frequency unpred: Cohen’s d 3.55, 2.42, 2.84, 2.54). There was a small, trend-level effect of right rs-Glx on the interaction between feature type and deviation (p=.05, F(1,16)=4.5, η2=0.06). Exploring this effect with a post-hoc regression analysis, including hearing accuracy and left rs-Glx as covariates, showed that the effect of right rs-Glx on unpred>pred difference was different in the two conditions, being a stronger positive relationship in duration (standardized beta = 0.49, t=2.15) and a weaker negative relationship in the frequency (standardized beta= −0.28, t=−1.04).

### The effect of rs-Glx on DMN and FPN between-network connectivity

We examined between-network connectivity, to test whether there was increased anticorrelation between DMN and FPN with increased rs-Glx. In all of four conditions (duration pred, duration unpred, frequency pred, frequency unpred), there was a small but significant positive connectivity between the networks (Cohen’s d: 0.81, 0.71, 0.84, 0.61). We did not find a significant unpred>pred contrast, or an effect of rs-Glx on the unpred>pred contrast. The only significant effect we did observe was a small main effect of condition (F(1,16)=5.2, p=.037, η^2^=0.04), due to higher positive connectivity between the networks during duration than frequency condition.

## Discussion

Glutamate has been related to coordinating the work of large brain networks during stimulus processing and cognitively demanding tasks. Glutamatergic receptors have been related to mediating the plasticity seen in the learning during oddball tasks: forming the model of the expected stimulation so that the deviance from the model could be detected. In this study we examined the effect of glutamate level in the primary auditory cortex on the brain activity during unpredictable auditory stimulation. We presented unpredictable blocks where the standard was repeatedly altered, which according to the predictive coding theories should lead to prediction error signaling. We observed that during the unpredictable blocks there was reduced connectivity within the auditory network and increased connectivity with prefrontal cortex, which is consistent with increased input from hierarchically higher levels during prediction error signaling. The rs-Glx level in the left auditory cortex was correlated to increased connectivity between the left auditory cortex and bilateral inferior parietal lobule. Further, we saw that in agreement with our hypotheses the rs-Glx level affected the coherence of both DMN and FPN during unpredictable stimulation: higher rs-Glx was associated with more coherence in FPN, whereas lower rs-Glx was associated with more coherence in DMN. However, this effect was specific to duration-deviant blocks, and depended on the hemisphere where rs-Glx was measured. Contrary to our hypotheses we did not find an effect of rs-Glx on connectivity within the hypothesized frontotemporal network based on source modeling of electrophysiological studies.

The areas in the inferior parietal lobe which showed connectivity with the left HG depending on the glutamate level were the areas in the caudal portion of the inferior parietal lobule (areas PGa and PGp). The rostral and caudal portion of IPL each exhibit a distinct cytoarchitectonic profile [19]. Also, these areas show functional segregation, with the rostral part (regions termed PF) related to motor and sensory functions, while the caudal part (PG) shows activation during complex cognitive abilities involved in language as well as attention [20]. The anatomical connectivity in the caudal areas is selectively oriented towards lateral occipital and temporal areas, following the inferior longitudinal fascicle; these regions are also connected to inferior frontal cortex, and show similarity in the distribution of glutamatergic and other receptors [20]. In particular, the caudal portion shows strong and preferential anatomical connectivity with areas in the temporal lobe, including the primary auditory cortex and planum temporale [21]. A significant left-right asymmetry in these connections has been found, with preferential connectivity between left primary auditory cortex and caudal IPL.

The involvement of IPL has been observed in several paradigms related to the current experiment. The IPL regions have been shown to be related to MMN for both duration- and frequency-deviance [8]. In the study by Molholm and colleagues, hemispheric differences were noted: the frequency-deviant blocks predominantly activated the right hemisphere, whereas the duration-related activation was bilateral or left-dominant. In our study, we did not find significant interaction effects suggesting the predominance of either hemisphere, however we did observe stronger effect in the right hemisphere, underlining its importance for both frequency- and duration-deviance. The important causal role for the parietal lobe in particular for duration-MMN has also been demonstrated by a TMS study showing reduced duration-MMN amplitude following a parietal TMS stimulation [22].

While there have been repeated attempts to relate the ventral attention network to oddball processing via the activations seen in the right inferior parietal region, this association deserves some careful anatomical consideration. The inferior portion of the region PGa (PGi), proposed based on the parcellation from the HCP project [23, 24], is suggested to be a part of the ventral attention network [25]. This region, however, appears to be more ventral than the areas found here; thus there is no clear evidence from this study of increased signaling between primary auditory cortex and ventral attention network during auditory deviance. Taking a more general view relative to the ventral attention network role of the IPL, Ptak has suggested that the role of the right IPL is related to shifting and maintaining of attention, interacting with a priority map which is held in the nearby inferior parietal sulcus [10].

Beyond attention, the parietal areas found here have been considered significant in representing prediction-related activity. During temporally unpredictable situations, right parietal activations have been observed, independent of WM load [26], and posterior parietal cortex has been associated with generating temporal expectancies [27]. In a recent meta-analysis, bilateral inferior parietal lobule (focus in IPS) was found to be sensitive to the magnitude of surprise in a reinforcement learning situation [28]. There appear to be fine-grained distinctions in the inferior parietal region, with the IPS being more sensitive to trial-specific surprise, whereas PGp is more activated for between-trial updating [29]. Intraparietal sulcus has been associated with relative uncertainty, which is maximal after a change in action-outcome mapping [30]. Also, uncertainty reduction via belief updating has been found to activate IPS together with superior frontolateral regions [31]. The present results suggest that the glutamatergic neurotransmission in sensory cortex plays an important role in the representation of expectancy, by influencing the connectivity between the expectancy-related brain areas and sensory brain regions.

Regarding the findings on the FPN and DMN, we predicted that across both deviance types there would be increased anticorrelation between the networks during unpredictable blocks, which should be stronger in participants with higher rs-Glx levels. We expected this anticorrelation to result from stronger connectivity within the FPN and weaker connectivity within DMN with higher rs-Glx levels. This hypothesis did not find support in the data. We did however find that during duration-deviant blocks there were effects of rs-Glx in the predicted direction on the within-network connectivity in both DMN as well as FPN. Here, the direction of the effects was as predicted: higher rs-Glx was related to higher FPN and lower DMN within-network connectivity during duration-deviant blocks, suggesting that individuals with higher rs-Glx mobilize their executive network during demanding perceptual environment, and correspondingly deactivate their DMN. However, the effects were dependent on the hemisphere where the rs-Glx was measured. In FPN, higher level of rs-Glx in the right hemisphere predicted higher within-network connectivity during the duration-deviant compared to duration-standard blocks. In DMN, higher levels of rs-Glx in the left hemisphere predicted lower within-network connectivity. Frontoparietal brain areas have been found to be engaged in estimating the environmental certainty, with increased environmental uncertainty leading to increased frontoparietal activation. Regularities, by contrast, have been shown to reduce metabolic demands in various areas. The dorsal frontoparietal network shows decreased activation for regular stimulation, where the prediction error is lower [32]. Also, predictability has been associated with greater DMN activity [33]. The importance of the right-sided rs-Glx may be related to different roles of the hemispheres for processing deviance in the stimulus features used here. Generally, representing certainty has been associated more with right than left frontoparietal network regions [34]. However, the frontoparietal areas in the right hemisphere in particular have been suggested to be important for duration discrimination [35]; also other studies have noted the particular relationship between parietal areas in the right hemisphere and duration-discrimination [36–38], also in brain-damaged patients [39, 40].

Taken together, our results show the importance of rs-Glx on the brain activation during unpredictable auditory stimulation. The primary auditory cortex connectivity with caudal portion of inferior parietal lobule (regions PGa and PGp and IPS) was higher in subjects with higher rs-Glx levels. This was observed independently of the deviating feature (duration or frequency). Rs-Glx did not have effect on the local connectivity within the temporal lobes. This agrees with previous findings of MRS-measured Glx: the effects are visible on long-range communication, not on local activation levels [17].

We suggest that the finding of increased communication between left HG and bilateral IPL reflects the increased calculations relevant to updating the representations of the auditory stimulation dependent on the surprise. Individuals with higher rs-Glx level in auditory cortex are more sensitive to the mismatch between predicted and actual stimulation as the increased glutamatergic neurotransmission allows more efficient signaling of prediction errors. This leads to more surprise-related activity in the hierarchically higher regions in the inferior parietal lobule.

We also demonstrate that the rs-Glx level has different effects on duration-deviance and frequency-deviance. Only the duration-deviant blocks were related to the network-wide activation as predicted, with higher rs-Glx related to increased connectivity within the FPN and decreased connectivity in DMN. The duration-representation has been shown to be particularly sensitive to conditions affecting the glutamatergic system, including high-risk to psychosis [41], and more sensitive to wider-spread calculations in the brain, including input from frontal sources [13]. The present results underline the difference in learning to predict different auditory features.

## Author contributions

Study design: KK, VV, AC. Data collection: VV, KK. Data analysis: KK, VV, AC, KS. Manuscript writing: KK, AC, KS.

## Availability of materials and data

The data is available from the corresponding author upon request.

