## Supplementary materials for "Glutamatergic modulation of auditory cortex connectivity with attentional brain networks in unpredictable perceptual environment"

*Supplementary figure 1*

The mean connectivity within each of the four conditions in a temporal-to-temporal connection (left HG to left pSTG) and temporal-to-frontal connection (left HG to left IFG).

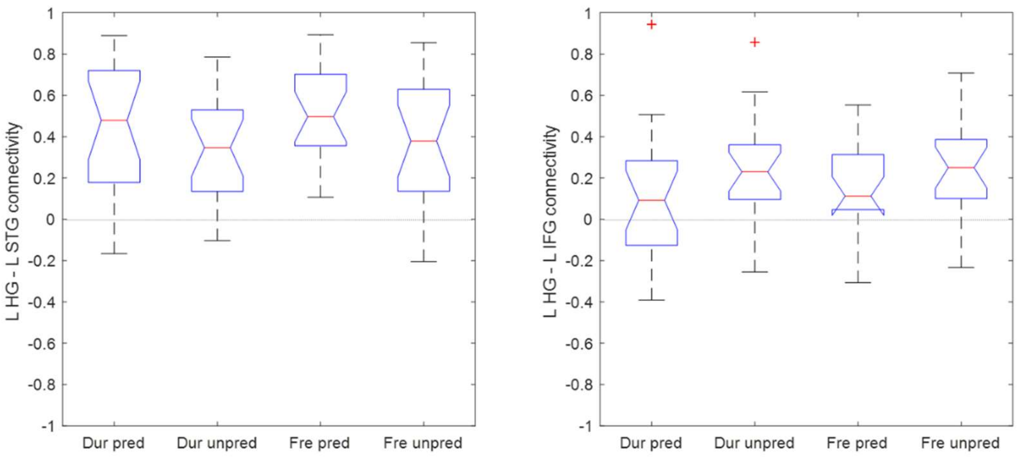

*Control analysis with water-scaled Glx and Cre*

To ensure that the effect of Glx/Cre on the connectivity between the auditory cortex and IPL reflects the contribution Glx, not interindividual differences in Cre levels, we performed a hierarchical regression analysis with the L_HG-R_IPL connectivity change in unpred>pred condition as the dependent variable. In the first step, Cre and hearing acuity were entered. The model was not significant (adjusted $R^2$ =.001, F(2,17)=1.01, p=.39). On the next step, Glx was entered; the resulting change in $R^2$ was significant ($R^2$ change .39, p=.003). The final model was significant (adjusted $R^2$=.405, F(3,16)=5.31, p=.01), thus the model including water-scaled Glx explained 41% of the variance in the L_HG-R_IPL connectivity in the unpred>pred contrast, a considerable improvement over the 0.1% variance explained in the model without Glx. In the final model (see Supplementary Table 1) both the Glx and Cre were significant predictors, with Cre having negative relationship. The hierarchical regression analysis with the L_HG-L_IPL connectivity change in unpred>pred showed similar outcome: adding Glx in the model led to significant increase in $R^2$, with the final model explaining almost half of the variance in the L_HG-L_IPL connectivity. As in the L_HG-R_IPL connection, the Cre had negative relationship with the connectivity. To explore this, we examined the relationship between Glx and Cre for multicollinearity. Glx and Cre were moderately correlated (Pearson r=.64). However, in the final model, the variance inflation factors (VIF) for Cre and Glx were small, suggesting that the contribution of both predictors on the dependent variable could be estimated without inflating the estimated regression coefficients. We note that when the portion of variance in Cre values shared between Glx and Cre has been accounted for, the remaining portion of interindividual variance in Cre is related to lower connectivity between the brain regions.

*Supplementary Table 1*

| **Variable/parameter** | **Model I stat (p)** | **Model II stat (p)** | **Model II VIF** |
|---|---|---|---|
| **L_HG-R_IPL** | | | |
| Hearing acuity | .18 (.45) | .31 (.12) | 1.09 |
| Cre | -.23 (.34) | -.73 (.006) | 1.68 |
| Glx | | 3.54 (.003) | 1.74 |
| Adjusted $R^2$ | .001 | .41 | |
| $R^2$ change | .11 (.39) | .39 (.003) | |
| **L_HG-L_IPL** | | | |
| Hearing acuity | .13 (.61) | .27 (.14) | 1.09 |
| Cre | -.18 (.47) | -.75 (.003) | 1.68 |
| Glx | | .94 (.001) | 1.74 |
| Adjusted $R^2$ | -.05 | .49 | |
| $R^2$ change | .06 | .51 | |
